## Supplemental Figures for "PASTA kinase-dependent control of peptidoglycan synthesis via ReoM is required for cell wall stress responses, cytosolic survival, and virulence in *Listeria monocytogenes*"

**Supplemental Figure 1.**

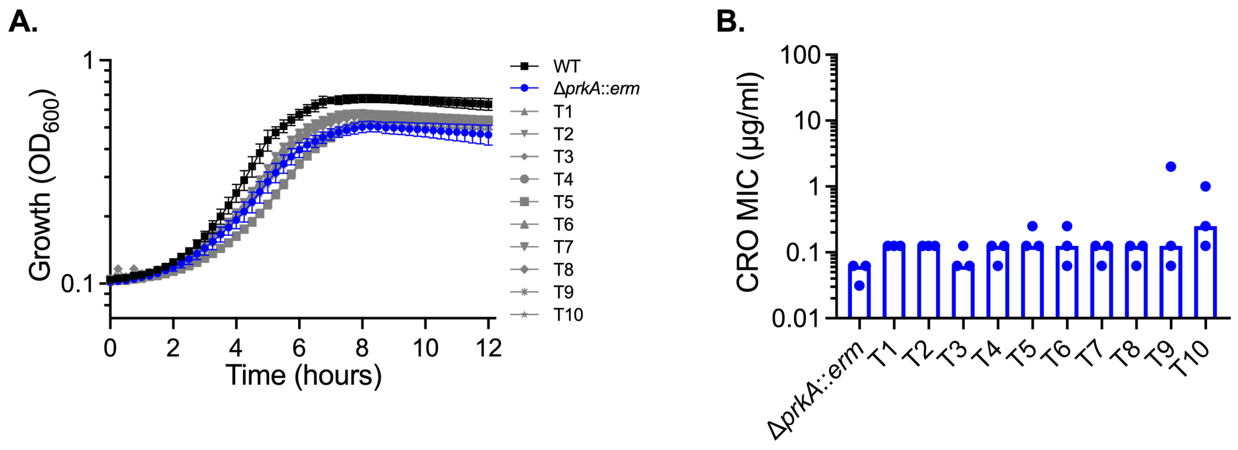


**Transductants of the Δ*prkA*::*erm* allele are not suppressors.** (**A**) Growth of WT, the Δ*prkA*::*erm* strain, and 10 Δ*prkA*::*erm* transductants (T1-T10) in BHI was monitored by $OD_{600}$. (**B**) Bars indicate median MICs of CRO for the indicated strains; n=3. No statistical differences between strains were found between the transductants and Δ*prkA*::*erm* by one-way ANOVA with Tukey's multiple comparisons test.

**Supplemental Figure 2.**

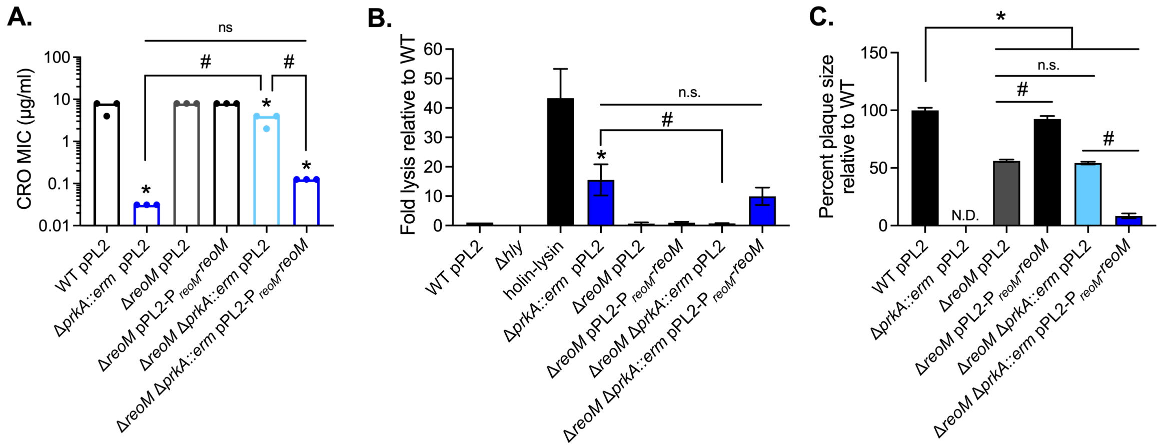


***Ex vivo* phenotypes of a Δ*reoM* mutant can be complemented by reintroduction of *reoM* controlled by its native promoter.** (**A**) Bars indicate median MICs of CRO for the indicated *L. monocytogenes* strains; n=3. (**B**) Intracellular bacteriolysis in immortalized *Ifnar*$^{-/-}$ macrophages. Macrophages were infected with the indicated strains carrying the pBHE573 reporter vector at an MOI of 10, and luciferase activity was measured 6 hours post-infection. Error bars indicate SEM; n=5. (**C**) Plaque formation in immortalized murine fibroblasts (L2 cells). L2s were infected with the indicated strains at an MOI of ~0.5, plaques were stained on day 4 of infection, and sizes were normalized to those of wild type. Error bars indicate SEM; data are averaged from a minimum of 64 plaques from three biological replicates. N.D., not detected. (**A-C**) *, $P < 0.05$ compared to wild type, and #, $P < 0.05$ for the indicated comparisons, by one-way ANOVA with Tukey's multiple comparisons test.

**Supplemental Figure 3.**

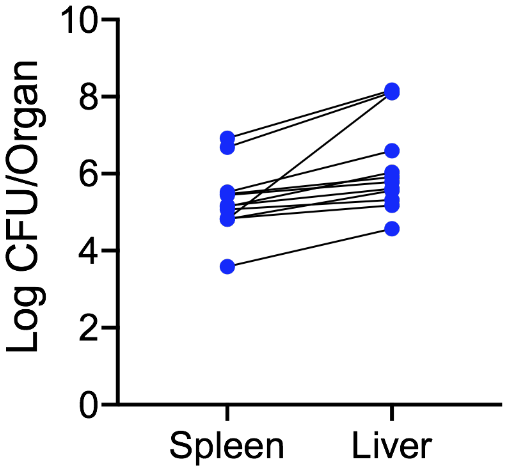


**Bacterial burdens from *in vivo* suppressor screen.** CFU were enumerated 72 hours post-infection with the EMS-mutagenized library of the Δ*prkA*::*erm* mutant.

**Supplemental Table 1.** All phosphopeptides identified in wild type and the Δ*prkA*$_{\text{cond}}$ mutant of *L. monocytogenes*, and PrkA-dependent phosphosites.

**Supplemental Table 2.** All phosphopeptides identified in wild type and the Δ*prkA*::*erm* mutant of *L. monocytogenes*.

**Supplemental Table 3.** Serine and threonine phosphopeptides with XCorr scores > 2.0 identified in wild type and the Δ*prkA*::*erm* mutant of *L. monocytogenes*.
